# No cell is an island: circulating T cell:monocyte complexes are markers of immune perturbations

**DOI:** 10.1101/411553

**Authors:** Julie G. Burel, Mikhail Pomaznoy, Cecilia S. Lindestam Arlehamn, Daniela Weiskopf, Ricardo da Silva Antunes, Yunmin Jung, Mariana Babor, Veronique Schulten, Grégory Seumois, Jason A. Greenbaum, Sunil Premawansa, Gayani Premawansa, Ananda Wijewickrama, Dhammika Vidanagama, Bandu Gunasena, Rashmi Tippalagama, Aruna D. deSilva, Robert H. Gilman, Mayuko Saito, Randy Taplitz, Klaus Ley, Pandurangan Vijayanand, Alessandro Sette, Bjoern Peters

## Abstract

Our results highlight for the first time that a significant proportion of cell doublets in flow cytometry, previously believed to be the result of technical artefacts and thus ignored in data acquisition and analysis, are the result of true biological interaction between immune cells. In particular, we show that cell:cell doublets pairing a T cell and a monocyte can be directly isolated from human blood, and high resolution microscopy shows polarized distribution of LFA1/ICAM1 in many doublets, suggesting *in vivo* formation. Intriguingly, T cell:monocyte complex frequency and phenotype fluctuate with the onset of immune perturbations such as infection or immunization, reflecting expected polarization of immune responses. Overall these data suggest that cell doublets reflecting T cell-monocyte *in vivo* immune interactions can be detected in human blood and that the common approach in flow cytometry to avoid studying cell:cell complexes should be revisited.

## Introduction

Communication between immune cells is a major component of immune responses, either directly through cell-cell contacts or indirectly through the secretion of messenger molecules such as cytokines. In particular, the physical interaction between T cells and antigen-presenting cells (APCs) is critical for the initiation of immune responses. APCs such as monocytes can take up debris from the extracellular environment, and will display fragments of it on their surface to T cells, which can identify potentially harmful, non-self antigens. There is paucity of data regarding T cell-APCs interactions in humans *in vivo*, but they appear to be highly diverse in terms of structure, length and function, depending on the nature and degree of maturation of the T cell and APC (Friedl & Storim, 2004).

Despite the importance of interactions between immune cells, many experimental techniques in immunology specifically avoid studying cell:cell complexes. The most notable example for this is in flow cytometry, in which cells are labeled with a panel of fluorochrome-conjugated antibodies, and each cell is then individually hit by a laser and its corresponding fluorescence emission spectra recorded. In this process, doublets (a pair of two cells) are routinely observed but are believed to be the results of technical artefacts due to *ex vivo* sample manipulation and are thus usually discarded, or ignored in data analysis (Kudernatsch, Letsch, Stachelscheid, Volk, & Scheibenbogen, 2013).

Blood is the most readily accessible sample in humans with high immune cell content. We and others have shown circulating immune cells contain critical information that can be used for diagnostic-, prognostic-and mechanistic understanding of a given disease or immune perturbation (Bongen, Vallania, Utz, & Khatri, 2018; Burel et al., 2018; Grifoni et al., 2018; Roy Chowdhury et al., 2018; Zak et al., 2016). Thus, whereas blood does not fully reflect what is occurring in tissues, it contains relevant immune information likely to be ‘leaking’ from the affected compartment. However, the presence of dual-cell complexes (and their content) has never been studied in the peripheral blood and in the context of immune perturbations. Monocytes are a subtype of phagocytes present in high abundance in the peripheral blood, which play a critical role in both innate and adaptive immunity (Jakubzick, Randolph, & Henson, 2017). In particular, monocytes have the capacity to differentiate into highly specialized APCs such as macrophages or myeloid DCs (Sprangers, de Vries, & Everts, 2016). More recently, it has been highlighted that they might directly function as APCs and thus contribute to adaptive immune responses (Jakubzick et al., 2017; Randolph, Jakubzick, & Qu, 2008).

We recently identified a gene signature in memory CD4+ T cells circulating in the peripheral blood that distinguishes individuals with latent TB infection (LTBI) from uninfected individuals (Burel et al., 2018). Surprisingly, this dataset also led to the discovery of a group of monocyte-associated genes co-expressed in memory CD4+ T cells whose expression is highly variable across individuals. We ultimately traced this discovery to a population of CD3+CD14+ cells that are not single cells but T cell:monocyte complexes present in the blood and that can be detected following immune perturbations such as disease or vaccination. The frequency and T cell phenotypes of these complexes appear to be associated with the nature of pathogen or vaccine. Thus, studying these complexes promises to provide insights into the impact of immune perturbation on APCs, T cells and their interactions.

## Results

### Unexpected detection of monocyte gene expression in CD4+ memory T cells from human subjects

We initially set out to investigate the inter-individual variability of gene expression within sorted memory CD4+ T cells from our previously characterized cohort of individuals with latent tuberculosis infection (LTBI) and uninfected controls (Burel et al., 2018). Within the 100 most variable genes, we identified a set of 22 genes that were highly co-expressed with each other (22-var set, **Figure 1 – source data 1**). Strikingly, many of the genes contained within the 22-var set were previously described as being highly expressed in classical monocytes (and to a lower extent non-classical monocytes) but not in T cells (**Figure 1B**, (Schmiedel et al., 2018)). In particular, the 22-var set contained the commonly used monocyte lineage marker CD14, the enzyme lysozyme LYZ and the S100 calcium binding proteins S100A8 and S100A9, which are known to be extremely abundant in monocytes (**Figure 1B**). By examining the flow cytometry data that were acquired during cell sorting and applying our memory CD4+ T cell gating strategy (**Figure 1 – figure supplement 1**), we identified that indeed there was a subpopulation within sorted memory CD4+ T cells that stained positive for CD14 (**Figure 1C**). More importantly, the proportion of memory CD4+ T cells that were CD14+ was positively correlated with the 22-var set expression (spearman correlation coefficient r=0.42, *p* < 0.0001, **Figure 1D**), suggesting that this cell subset is responsible for the expression of the monocyte-associated genes identified in Figure 1A. The CD14+ memory CD4+ T cell population has similar forward and side scatter (FSC/SSC) values to other memory CD4+ T cells and was thus sorted along with conventional CD14- memory CD4+ T cells (**Figure 1 – figure supplement 2**). In particular, there was no indication that CD14+ memory CD4+ T cells were the product of a technical artefact, such as dead cells or a compensation issue.

**Figure 1.**
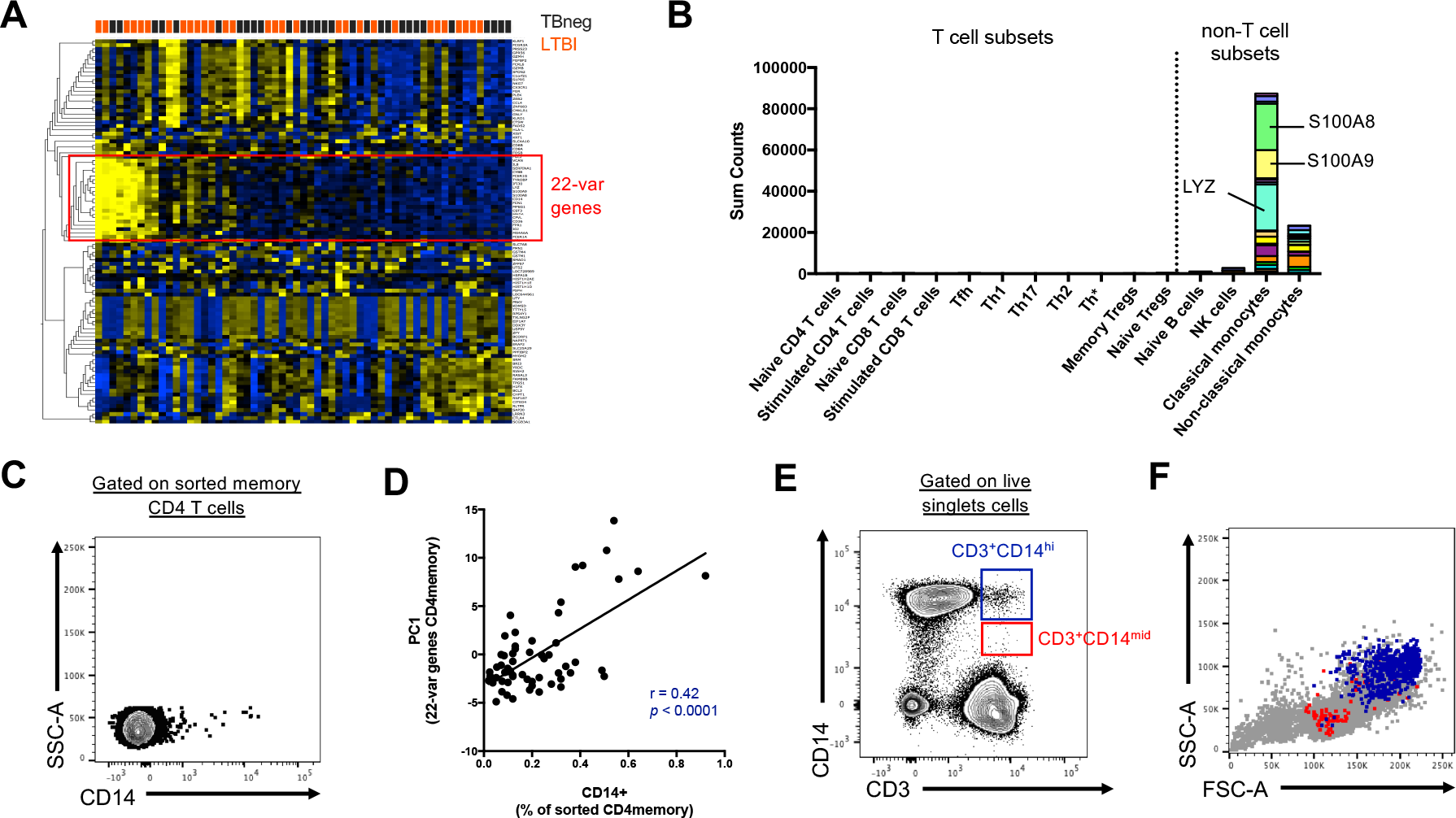
Two cell populations expressing both T cell (CD3) and monocyte (CD14) surface markers exist in the live singlet cell population of PBMC from human subjects. A) The top 100 most variable genes in memory CD4+ T cells across TB uninfected (TBneg) and LTBI infected subjects. B) Immune cell type specific expression of the 22-var genes identified in A). Every bar consists of stacked sub-bars showing the TPM normalized expression of every gene in corresponding cell type. Expression of genes for the blood cell types shown were taken from the DICE database ((Schmiedel et al., 2018), http://dice-database.org/). C) Detection of CD14+ events within sorted CD4+ memory T cells and D) non-parametric spearman correlation between their frequency and the PC1 from the 22-var genes. E) Gated on ‘singlet total live cells’, two populations of CD3+CD14+ cells can be identified based on the level of expression of CD14. F) Based on FSC and SSC parameters, CD3+CD14hi cells are contained within the monocyte gate, whereas CD3+CD14mid cells are contained within the lymphocyte gate. Data were derived from 30 LTBI subjects and 29 TB uninfected control subjects.

### Distinct CD3+CD14+ cell populations are present in the monocyte vs. the lymphocyte size gate

To further investigate the origin of the CD14+ T cell population, we analyzed our flow cytometry data, this time not restricting to the compartment of sorted memory T cells, but looking at all cells. When gating on live FSC/SSC (including both monocytes and lymphocytes) singlet cells, two populations of CD3+CD14+ could be readily identified: CD3+CD14hi cells and CD3+CD14mid cells (**Figure 1E**). CD3+CD14hi cells were predominantly contained within the monocyte size gate, whereas CD3+CD14mid cells were contained within the lymphocyte size gate (**Figure 1F**).

### CD3+CD14+ cells consist of T cells bound to monocytes or monocyte debris

To better understand the nature of CD3+CD14+ cells, we aimed to visualize the distribution of their markers using imaging flow cytometry. Live events were divided into monocytes (CD3-CD14+), T cells (CD3+CD14-), CD3+CD14hi cells, and CD3+CD14mid cells (**Figure 2A**), and a random gallery of images was captured for each population. As expected, monocytes and T cells contained exclusively single cells that expressed either CD14 (monocytes) or CD3 (T cells), respectively (**Figure 2B**, *first and second panel*). To our surprise, CD3+CD14hi cells contained predominantly two cells, sometimes even three cells, but no single cells (**Figure 2B**, *third panel*). The doublets (or triplets) always contained at least one CD14+ cell, and one CD3+ cell (**Figure 2B**, *third panel*). CD3+CD14mid cells contained predominantly single CD3+ cells, but also some doublets of one CD3+ cell and one CD14+ cell, but with CD14 expression lower than average monocytes (**Figure 2B**, *fourth panel*). The majority of CD3+ T cell singlets in the CD3+CD14mid population, but not in the CD3+CD14-T cell population, contained CD14+ particles, often seen at the periphery of the CD3+ T cell membrane (**Figure 2B-C**). Looking more closely at the CD14+ particles contained within the CD3+CD14mid population using confocal microscopy, they were found to have size and shape similar to cell debris (**Figure 2D**). To confirm our initial observation, we repeated the experiment with multiple individuals, and compared for each cell population the aspect ratio and area from the brightfield parameter collected with the image stream. Doublets are known to present a larger area but reduced aspect ratio, when compared to single cells. Thus, their overall ratio (area vs aspect ratio) is greater than in single cells. As expected, the area vs aspect ratio was significantly higher for CD3+CD14hi cells and CD3+CD14mid cells compared to single monocytes and T cells, and events in these two cell populations were found predominantly in the ‘doublet gate’ (**Figure 2E-F**). CD3+CD14hi cells also had a significantly higher ratio compared to CD3+CD14mid cells (**Figure 2F**).

**Figure 2.**
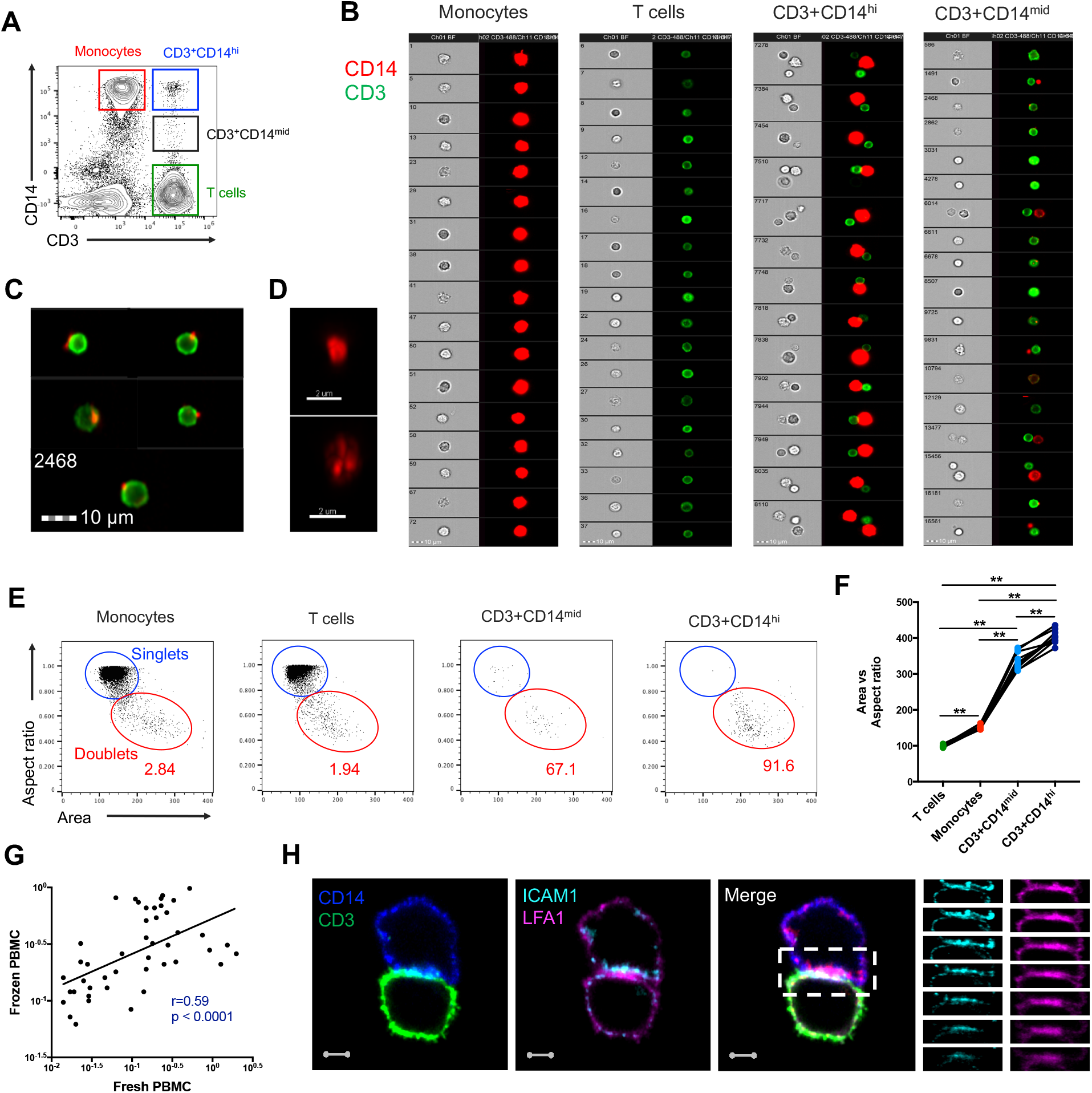
CD3+CD14+ cells are tightly bound T cell:monocyte complexes that represent *in vivo* association. A) Gating strategy and B) random gallery of events for monocytes (CD14+CD3-), T cells (CD3+CD14-), CD3+CD14hi cells and CD3+CD14mid cells determined by imaging flow cytometry (ImageStreamX, MkII Amnis Amnis). CD14+ cell debris were identified within CD3+CD14mid cells C) by imaging flow cytometry and D) confocal microscopy after bulk population cell sorting. E) Plots and F) Ratio of Aspect ratio vs Area of the brightfield parameter for monocytes (CD14+CD3-), T cells (CD3+CD14-), CD3+CD14hi cells and CD3+CD14mid cells, determined by imaging flow cytometry. G) Non-parametric Spearman correlation of the frequency of live singlets CD3+CD14+ cells in paired fresh PBMC vs cryopreserved PBMC derived from 45 blood draws of healthy subjects. H) Single z-plan (0µm) images (*left*) and z-plane stacks (*right*) of the region marked (dashed rectangle) from one sorted CD3+CD14+ T cell:monocyte complex displaying accumulation of LFA1 and ICAM1 at the interface. Images show expression of CD14 (blue), CD3 (green), ICAM1(Cyan), and LFA1 (Magenta). Relative z-positions are indicated on the right, and scale bars represent 2 µm. Imaging flow cytometry data was derived from 10 subjects across three independent experiments and microscopy data was representative of the analysis of n=105 CD3+CD14+ complexes isolated from 3 subjects across three independent experiments.

Taken together, these results demonstrate that CD3+CD14hi cells are tightly bound T cell:monocyte complexes, in such strong interaction that sample processing and flow cytometry acquisition did not break them apart. Conversely, the CD3+CD14mid population appears to predominantly consist of single CD3+ T cells with attached CD14+ cell debris. This conclusion is further supported by CD3+CD14hi complexes being found in the monocyte size gate, whereas CD3+CD14mid cells were falling into the lymphocyte size gate (**Figure 1F**).

### T cell:monocyte complexes are not the result of cryopreservation or PBMC isolation

Next, we sought to determine whether the physical association of T cells and monocytes within the T cell:monocyte complexes was the result of random cellular proximity during *ex vivo* sample manipulation, or if the complexes are originally present in peripheral blood. We could readily detect T cell:monocyte complexes in freshly isolated PBMC, and at similar frequencies as the same samples after cryopreservation (**Figure 2G**). In another set of samples, using red blood cell (RBC) magnetic depletion (and thus minimal sample manipulation), we could successfully identify T cell:monocyte complexes directly from whole blood at frequencies matching the same sample after PBMC isolation (**Figure 2 - figure supplement 1**). Taken together these data rule out that the PBMC sample preparation or cryopreservation could be responsible for T cell:monocyte complexes formation and thus suggest their presence *in vivo* in peripheral blood.

### T cell:monocyte complexes show increased expression of adhesion molecules at their interface

During T cell recognition of epitopes on APCs such as monocytes, the two cells are known to form an ‘immune synapse’ at their contact point, which is stabilized by key adhesion molecules such as LFA1 on the T cell, and ICAM1 on the APC (Dustin, 2014). Upon interaction these two molecules undergo a drastic redistribution by focusing almost exclusively at the cell:cell point of contact, thus forming a ‘ring’ that can be visualized (Wabnitz & Samstag, 2016). To identify candidate immunological synapses in T cell:monocyte complexes, we used high resolution Airyscan images of sorted doublets (see **Figure 2 – figure supplement 2** for sorting strategy). Almost a third (thirty out of 105, 29%) of doublets analyzed from three different individuals displayed accumulation and polarization of ICAM1 and LFA1 at their interfaces (**Figure 2H**). The percentage of polarized doublets ranged from 17 to 67% between the subjects. In seven doublets, CD3 also accumulated together with LFA1 (**Figure 2 – figure supplement 3**). However, we did not find welldeveloped, classical immunological synapses, defined by central accumulation of CD3 and LFA1 exclusion from central region of a synapse (Monks, Freiberg, Kupfer, Sciaky, & Kupfer, 1998; Thauland & Parker, 2010). Overall, this suggests that a significant fraction of the detected T cell:monocyte complexes utilizes adhesion markers associated with T cell:APC synapse formation to stabilize their interaction, but they do not appear to being currently undergoing active TCR signaling.

### The frequency of T cell:monocyte complexes varies in the context of diverse immune perturbations

Next, we thought to examine whether the formation of T cell:monocyte complexes is dysregulated following immune perturbations. In order to accurately assess and compare the frequency of complexes between cells of different types across different donor cohorts, we need to take into account that their frequency is dependent on the abundance of its two components. Indeed, in healthy subjects, where we expect constant affinity between T cells and monocytes, we observed that the frequency of CD3+CD14+ cells is a linear function of the product of singlet monocyte and T cell frequencies (**Figure 3A**). To correct for this, we elected to express the abundance of T cell:monocytes complexes as a constant of association Ka, where similarly to a constant of chemical complex association, the frequency of T cell:monocyte complexes is divided by the product of the frequency of both T cells and monocytes (**Figure 3B**). As T cell and monocyte frequencies in the blood can fluctuate greatly during immune perturbations, the Ka is a more accurate readout of the likelihood of T cell:monocyte complex formation as opposed to raw frequencies.

**Figure 3.**
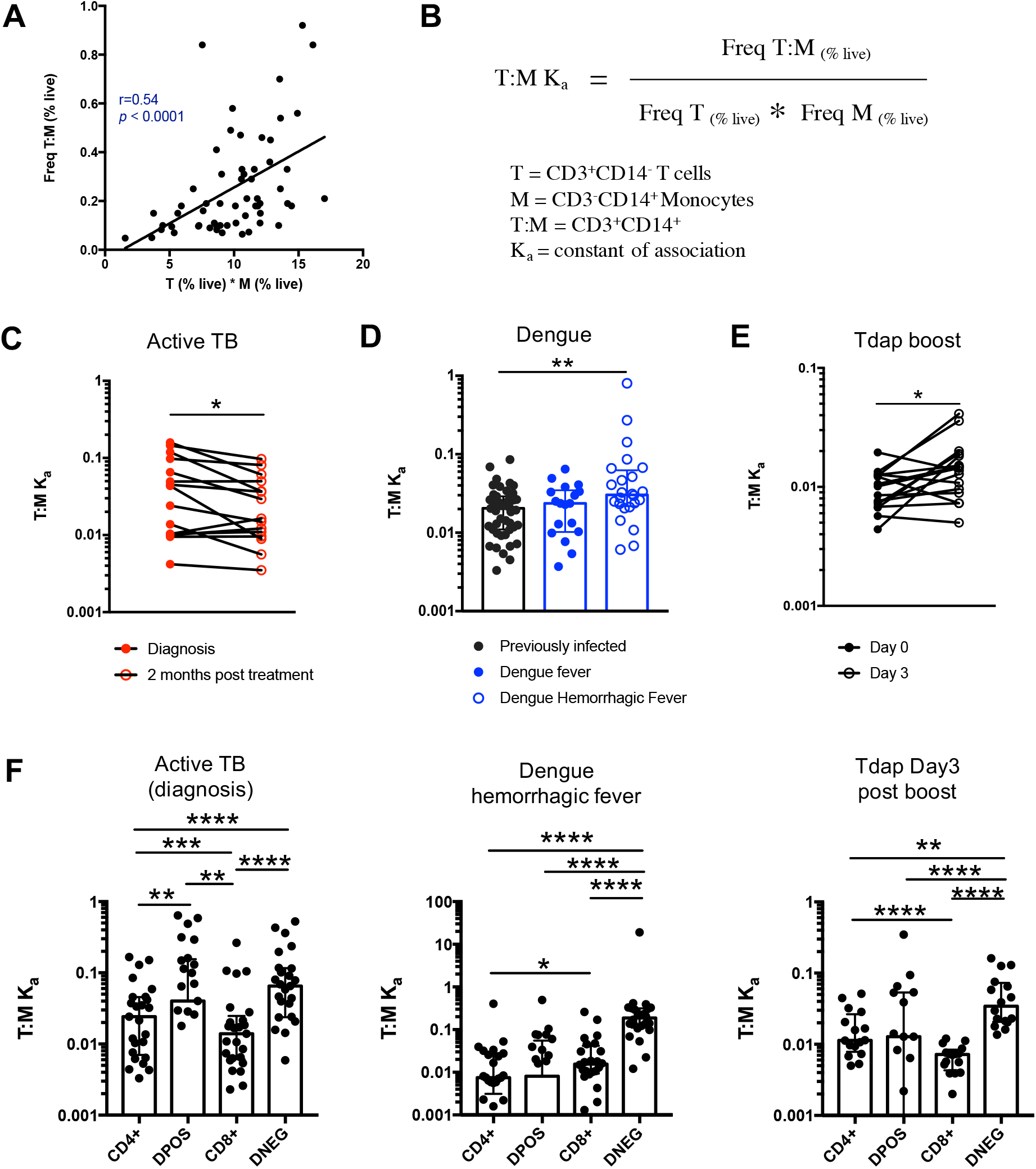
The constant of association Ka between monocytes and T cells (and T cell subsets) varies with the presence and nature of immune perturbations. A) Non-parametric spearman correlation between the frequency of T cell:monocyte complexes and the product of singlet T cells and monocyte frequencies in healthy subjects (n=59). B) Formula for the calculation of the T cell:monocyte constant of association Ka. T cell:monocyte complexes constant of association Ka in C) active TB subjects at diagnosis and 2 months post treatment (n=15), D) individuals with acute dengue fever (n=18), acute dengue hemorrhagic fever (n=24) or previously infected (n=47) and E) previously vaccinated healthy adults (n=16) before and three days post boost with Tdap vaccine, calculated as explained in B). F) The constant of association Ka between monocytes and T cell subsets in active TB subjects at diagnosis (n=25), individuals with acute dengue hemorrhagic fever (n=24) and previously vaccinated healthy adults three days post boost with Tdap vaccine (n=16), calculated as explained in B). Statistical differences over time and across cell populations within subjects were determined using the non-parametric paired Wilcoxon test; other statistical differences were determined using the non-parametric Mann-Whitney test; *, *p* < 0.05; **, *p* < 0.01; ***, *p* < 0.01; ****, *p* < 0.0001. Plots represent individual data points, median and interquartile range across all subjects within each cohort. Raw frequencies of T cell:monocyte complexes for the different disease cohorts are available on Figure 3 – figure supplement 3.

We first investigated the T cell:monocyte Ka in the context of two diseases where monocytes are known to be important, namely active tuberculosis (TB) infection and dengue fever. In the case of TB, although macrophages are known to be the primary target for *Mycobacterium tuberculosis* (Mtb) infection and replication, monocytes can also be infected and contribute to the inflammatory response (Srivastava, Ernst, & Desvignes, 2014). In active TB subjects, we found a significant decrease in T cell:monocyte Ka at 2 months post treatment (**Figure 3C**). At the time of diagnosis, some subjects displayed a Ka much higher than any uninfected or LTBI individuals, but because of the high heterogeneity within the active TB cohort, these differences did not reach statistical significance (**Figure 3 – figure supplement 1**). Dengue virus predominantly infects monocytes in the peripheral blood (Kou et al., 2008), and circulating monocyte infection and activation is increased in dengue hemorrhagic fever (the more severe form of dengue fever) (Durbin et al., 2008). In subjects with acute dengue fever from Sri Lanka, patients that developed hemorrhagic fever had higher T cell:monocyte Ka upon hospitalization compared to healthy, previously infected subjects (blood bank donors seropositive for dengue antibodies) (**Figure 3D**). In contrast, patients with a less severe form of acute dengue infection showed no significant difference in T cell:monocyte Ka compared to healthy, previously infected donors (**Figure 3D**).

To assess whether vaccination also impacted the formation of T cell:monocyte complexes, we obtained samples from healthy adults that received the tetanus, diphtheria and pertussis (Tdap) booster vaccination. We indeed observed a significantly higher T cell:monocyte Ka at three days post boost compared to baseline (**Figure 3E**), but no significant changes at one, seven or fourteen days post boost (**Figure 3 – figure supplement 2**). Taken together, these data confirm that circulating T cell:monocyte complexes can be found directly *ex vivo* in different immune perturbations, and their likelihood of formation is associated with clinical parameters such as disease severity, and they fluctuate as a function of time post treatment and post vaccination.

### T cells with different phenotypes are found in T cell:monocyte complexes dependent on the nature of the immune perturbation

Finally, we reasoned that if immune perturbations increase the formation of T cell:monocyte complexes, then the nature of the T cells contained in the complexes could provide insights into which T cells are actively communicating with monocytes *in vivo*. In particular, the T cell subsets that will associate with an APC for the different perturbations studied above are expected to be distinct, and thus their likelihood to form a complex with a monocyte might differ too. The Tdap vaccine contains exclusively protein antigens and is known to elicit predominantly memory CD4+ T cell responses (da Silva Antunes et al., 2018). *Mtb* is a bacterial pathogen known to trigger strong CD4+ responses (Lindestam Arlehamn et al., 2016) as opposed to dengue virus, which is a viral antigen and thus expected to elicit CD8+ responses.

Similarly to global T cell:monocyte complexes (Figs. 3C-E), we calculated for each CD4/CD8 T cell subset its constant of association Ka with monocytes. In subjects with active TB, the Ka between monocytes and CD4+CD8+ (DPOS) T cells or CD4+ T cells was significantly higher than for CD8+ T cells (**Figure 3F**) and both DPOS and CD4+ T cell:monocyte complexes had higher Ka in active TB compared to dengue hemorrhagic fever (**Figure 3 – figure supplement 3**). Dengue hemorrhagic fever showed a higher T cell:monocyte Ka for CD8+ over CD4+ cells whereas Tdap day 3 post boost showed the opposite, with highest Ka for CD4+ over CD8+ cells (**Figure 3F**). The CD8+ T cell:monocyte Ka was also higher in Dengue and active TB compared to Tdap boost (**Figure 3 – figure supplement 3**). Thus, the magnitude of Ka in CD4+ vs CD8+ T cell subsets matched what is expected based on the nature of immune perturbation. Interestingly, for all three immune perturbations studied the highest Ka with monocytes across all T cell subsets was for CD4CD8 (DNEG) T cells (**Figure 3D**), and this effect was most pronounced in dengue (**Figure 3 – figure supplement 3**). These cells could constitute gamma-delta T cells that are known to be strongly activated in the peripheral blood during acute dengue fever (Tsai et al., 2015).

In summary, these data indicate that the T cell subsets that are preferentially associated with monocytes differ from their individual frequencies in PBMC, and follow different patterns in the three systems studied, further supporting the notion that these complexes are not the result of random association, and are specific to the nature of the immune perturbation.

## Discussion

The unexpected detection of monocyte genes expressed in cells sorted for memory T cell markers led to the discovery that a population of CD3+CD14+ cells exist within the ‘live singlet’ events gate and that these cells are T cells that are tightly associated with monocytes, and less frequently, with monocyte-derived debris. Their presence in freshly isolated cells and the fact that a significant fraction of the complexes showed enriched expression for LFA1/ICAM1 adhesion molecules at their interface, suggest that they are not the product of random association of cells during processing, but represent true interactions that occurred *in vivo* prior to the blood draw. The frequency of T cell:monocyte complexes fluctuated over time in the onset of immune perturbations such as following TB treatment or Tdap boost immunization and correlated with clinical parameters such as disease severity in the case of dengue fever. Furthermore, the T cell subset in preferential association within the monocyte in a complex varies in function of the nature of the immune perturbation.

Thus, circulating CD3+CD14+ complexes appear to be the result of *in vivo* interaction between T cells and monocytes. Because cells are in constant motion in the bloodstream, it is possible that T cell:monocyte complex formation does not initially occur in peripheral blood. While we cannot exclude that this is the case, we consider it more likely that their formation occurs in tissues or draining lymph nodes, and the complexes are then ‘leaking’ into the peripheral circulation. The most studied physical interaction between T cells and monocytes is the formation of immune synapses. We found that about a third of complexes displayed LFA1/ICAM1 mediated interaction similarly to immune synapses, but no CD3 polarization. The immune synapse formation is a highly diverse event in terms of length and structure (Friedl & Storim, 2004), so it is possible that not all detected complexes are at the same stage in the interaction. In some complexes, the nature (and structure) of the architectural molecules forming the cell:cell contact might differ from traditional immune synapses, too. Studying the nature and physical properties of these interactions could provide insights into how T cells and monocytes can physically interact. Additionally, because monocytes are not the only cell type known to associate with T cells, we think the ability to form complexes with T cells should not be restricted to monocytes, but could apply more broadly to any APC. Thus, it is likely that other types of complexes pairing a T cell and other APCs such as B cells or dendritic cells can be found in the peripheral blood.

Increased immune cell:cell interactions might not necessarily always correlate with onset of immune perturbations. Nevertheless, our preliminary data suggest that determining the constant of association Ka of the T cell:monocyte (and likely more broadly any T cell:APC) complexes can indicate the presence of an immune perturbation to both clinicians and immunologists. In dengue infected subjects, a higher T cell:monocyte Ka at time of admission was associated with dengue hemorrhagic fever, the more severe form of disease. The distinction between hemorrhagic vs. non-hemorrhagic fever may become clear only days into hospitalization, so the ability to discriminate these two groups of individuals at the time of admission has potential diagnostic value. In the case of active TB, subjects presented a very high variability at diagnosis that might reflect the diverse spectrum associated with the disease (Pai et al., 2016), but for all subjects a significant decrease in T cell:monocyte Ka was observed upon treatment. This could thus be a tool to monitor treatment success and predict potential relapses. It will of course be necessary to run prospective trials to irrefutably demonstrate that the likelihood of association between T cells and monocytes have predictive power with regard to dengue disease severity or over the course of TB treatment. Additionally, the T cell:monocyte Ka was increased three days following Tdap booster vaccination. Therefore, in vaccine trials, it could be examined as an early readout to gauge how well the immune system has responded to the vaccine. Finally, in apparently ‘healthy’ populations, or those with diffuse symptoms, an unusually high T cell:monocyte Ka in an individual could be used as an indicator of a yet to be determined immune perturbation.

Beyond detecting abnormal frequencies of T cell:monocyte complexes, characterizing the T cells and monocytes in these complexes might provide insights into the nature of immune perturbation and subsequent immune response based on which complexes were formed. Our data suggest that there are drastic differences in terms of T cell subsets in the complexes. Despite their lower frequency over CD4+ and CD8+ T cells in the peripheral blood, DNEG T cells show a clear increased association with monocytes. Gamma-delta T cells constitute the majority of circulating DNEG T cells in humans, and LFA1 dependent crosstalk between gamma-delta T cells and monocytes has been shown to be important in the context of bacterial infections (Eberl et al., 2009), which might be also generalized to viral infections. Thus, the DNEG T cell:monocyte complexes might well represent a novel type of interaction between T cells and monocytes, not necessarily involving classical alpha-beta T cells or involving the formation of ‘traditional’ immune synapses. Aside from the enrichment for DNEG cells in T cell:monocyte complexes in all samples analyzed, we also observed that CD4 vs CD8 phenotype of the T cell present in complexes depends on the nature of the immune perturbation studied, and reflects the expected polarization of immune responses. Thus, looking for additional characteristics from T cells and monocytes present in the complexes, such as the expression of tissue homing markers, specific TCRs and their transcriptomic profile might provide further information about the fundamental mechanisms underlying immune responses to a specific perturbation.

Why were T cell:monocyte complexes not detected and excluded in flow cytometry based on gating strategies to avoid doublets? Surprisingly, all usual parameters (pulse Area (A), Height (H) and Width (W) from forward and side scatter) looked identical between T cell:monocyte complexes and singlet T cells or monocytes. The only parameter that could readily distinguish between intact CD3+CD14hi complexes and single T cells or monocytes was the brightfield area parameter from the imaging flow cytometer, which is a feature absent in non-imaging flow cytometry. Thus, it seems that gating approaches and parameters available in conventional flow cytometry are not sufficient to completely discriminate tightly bound cell pairs from individual cells.

Given that T cell:monocyte complexes are not excluded by conventional FACS gating strategies, why were they not reported previously? Examining our own past studies, a major reason is that lineage markers for T cells (CD3), B cells (CD19) and monocytes (CD14) are routinely used to remove cells not of interest in a given experiment by adding them to a ‘dump channel’. For example, most of our CD4+ T cell studies have CD8, CD19 and CD14, and dead cell markers combined in the same channel (Arlehamn et al., 2014; Burel et al., 2018). Other groups studying e.g. CD14+ monocytes are likely to add CD3 to their dump channel. This means that complexes of cells that have two conflicting lineage markers such as CD3 and CD14 will often be removed from datasets early in the gating strategy. Additionally, the detection of complexes by flow cytometry is not straightforward. In our hands, we have found that conventional flow analyzers give low frequency of complexes and poor reproducibility in repeat runs. This is opposed to cell sorters, presumably due to differences in their fluidics systems, which puts less stress on cells and does not disrupt complexes as much. Both the routine exclusion of cell populations positive for two conflicting lineage markers and the challenges to reproduce such cell populations on different platforms has likely contributed to them not being reported.

Moreover, even if a panel allows for the detection of complexes, and there is a stable assay used to show their presence, there is an assumption in the field that detection of complexes is a result of experimental artefacts. For example, we found a report of double positive CD3+CD34+ cells detected by flow cytometry in human bone marrow, which followed up this finding and found them to be doublets using microscopy imaging. The authors concluded that these complexes are the product of random association and should be ignored (Kudernatsch et al., 2013). Their conclusion may well be true for their study, but it highlights a common conception in the field of cytometry that pairs of cells have to be artefacts. Another study described CD3+CD20+ singlets cells observed by flow cytometry as doublets of T cells and B cells, and also concluded them to be a technical artefact, in the sense that these cells are not singlets double expressing CD3 and CD20 (Henry et al., 2010). In this case however, authors pointed out that ‘Whether the formation of these doublets is an artefact occurring during staining or is a physiologic process remains to be determined’ (Henry et al., 2010).

We ourselves assumed for a long time that we might have an artefact finding, but given the persistent association of T cell:monocyte complexes frequency and phenotype with clinically and physiologically relevant parameters, we came to a new conclusion: cells are meant to interact with other cells. Thus, detecting and characterizing complexes of cells isolated from tissues and bodily fluids, can provide powerful insights into cell:cell communication events that are missed when studying cells as singlets only.

## Material and methods

### Ethics statement

Samples from TB uninfected individuals were obtained from the University of California, San Diego Antiviral Research Center clinic (AVRC at UCSD, San Diego) and National Blood Center (NBC), Ministry of Health, Colombo, Sri Lanka, in an anonymous fashion as previously described (Burel et al., 2017). Samples from individuals with LTBI were obtained from AVRC at UCSD, San Diego, and the Universidad Peruana Cayetano Heredia (UPCH, Peru). Longitudinal active TB samples were obtained from National Hospital for Respiratory Diseases (NHRD), Welisara, Sri Lanka. Dengue previously infected samples were obtained from healthy adult blood donors from the National Blood Center (NBC), Ministry of Health, Colombo, Sri Lanka, in an anonymous fashion as previously described (Weiskopf et al., 2013). Acute dengue fever samples were collected at National Institute of Infectious Diseases, Gothatuwa, Angoda, Sri Lanka and the North Colombo Teaching Hospital, Ragama, in Colombo, Sri Lanka. Longitudinal Tdap booster vaccination samples were obtained from healthy adults from San Diego, USA. Ethical approval to carry out this work is maintained through the La Jolla Institute for Allergy and Immunology Institutional Review Board, the Medical Faculty of the University of Colombo (which served as a National Institutes of Health–approved institutional review board for Genetech) and the John’s Hopkins School of Public Health Institutional Review Board (RHG holds dual appointment at UPCH and JHU). All clinical investigations have been conducted according to the principles expressed in the Declaration of Helsinki. All participants, except anonymously recruited blood bank donors in Sri Lanka, provided written informed consent prior to participation in the study.

### Subjects and samples

LTBI status was confirmed in subjects by a positive IFN-γ release assay (IGRA) (QuantiFERON-TB Gold In-Tube, Cellestis or T-SPOT.TB, Oxford Immunotec) and the absence of clinical and radiographic signs of active TB. TB uninfected control subjects were confirmed as IGRA negative. Active Pulmonary TB was defined as those exhibiting symptoms of TB, and are positive by sputum and culture as confirmed by the National Tuberculosis Reference Laboratory (NTRL, Welisara, Sri Lanka). Sputum was further confirmed positive for TB by PCR at Genetech (Sri Lanka). Active TB patients in this study were confirmed negative for HIV, HBV and HCV. Upon enrollment within seven days of starting their anti-TB treatment, active TB patients provided their first blood sample, followed by a second blood sample two months after initial diagnosis. Acute dengue fever and previously infected samples were classified by detection of virus (PCR+) and/or dengue-specific IgM and IgG in the serum. Laboratory parameters such as platelet and leukocyte counts, hematocrit, hemoglobulin, AST, ALT and if applicable an ultrasound examination of the chest and abdomen or an X-ray were used to further diagnose patients with either dengue fever (DF) or dengue hemorrhagic fever (DHF), a more severe form of disease, according to WHO’s guidelines. Longitudinal Tdap booster vaccination samples were obtained from individuals vaccinated in childhood, and boosted with the DTP vaccine Tdap (Adacel). Blood samples were collected prior, one day, three days, seven days and fourteen days post boost. For all cohorts, PBMC were obtained by density gradient centrifugation (Ficoll-Hypaque, Amersham Biosciences) from leukapheresis or whole blood samples, according to the manufacturer’s instructions. Cells were resuspended to 10 to 50 million cells per mL in FBS (Gemini Bio-Products) containing 10% dimethyl sulfoxide (Sigma) and cryopreserved in liquid nitrogen.

### Flow cytometry

Surface staining of fresh or frozen PBMC was performed as previously described in (Burel et al., 2017). Briefly, cells were stained with fixable viability dye eFluor506 (eBiosciences) and various combinations of the antibodies listed in **Table supplement 1** for 20min at room temperature. Acquisition was performed on a BD LSR-I cell analyzer (BD Biosciences) or on a BD FACSAria III cell sorter (BD Biosciences). Compensation was realized with single-stained beads (UltraComp eBeads, eBiosciences) in PBS using the same antibody dilution as for the cell staining.

### Imaging flow cytometry

For the visualization of CD3+CD14+ cells, frozen PBMC were thawed and stained with CD3-AF488 and CD14-PE or CD14-AF647 (see Table supplement 1 for antibody details) as described in the flow cytometry section above. After two washes in PBS, cells were resuspended to 10×10^6^ cells/mL in FACS buffer containing 5μg/mL Hoechst (Invitrogen) and 1μg/mL 7-AAD (Biolegend) and stored at 4°C protected from light until acquisition. Acquisition was performed with ImageStreamX MkII (Amnis) and INSPIRE software version 200.1.620.0 at 40X magnification and the lowest speed setting. A minimum of 4,000 CD3+CD14+ events in focus were collected. Data analysis was performed using IDEAS version 6.2.183.0.

### Sample preparation for microscopy

For the visualization of LFA1/ICAM1 polarization on T cell:monocyte complexes, frozen PBMC were thawed and resuspended in blocking buffer (2% BSA, 10mM EGTA, 5mM EDTA, 0.05% Sodium Azide in 1X PBS) supplemented with 2ul of Trustain FcR blocking reagent (BioLegend) for 10min on ice. Antibodies (anti-human CD3-AF488, CD14-BV421, ICAM1-AF568, LFA1-CF633 or LFA1-AF647, see Table supplement 1 for antibody details) were added and incubated for 20 min on ice, and then washed twice with FACS buffer (PBS containing 0.5% FBS and 2mM EDTA, pH 8). Cells were fixed with 4% Paraformaldehyde, 0.4% Glutaldehyde, 10mM EGTA, 5mM EDTA, 0.05 Sodium Azide, 2% sucrose in PBS for 1 h on ice, and then washed twice with MACS buffer. Cells were resuspended in 0.5-1mL of MACS buffer, and kept at 4°C until sorting. Cell sorting was performed on a BD Aria III/Fusion cell sorter (BD Biosciences). CD3+CD14+, CD3+CD14-T cells and CD14+CD3- monocytes were sorted (see gating strategy **Figure supplement 1B**) and each separately plated on a well of a µ-Slide 8 Well Glass Bottom chamber (Ibidi) that was freshly coated with poly-L-lysine (0.01%) for 30 min RT before use. For in-house antibody labeling, an Alexa Fluor^™^ 568 antibody labeling kit and a Mix-n-Stain^™^ CF®633 Dye antibody labeling kit (Sigma) were used according to manufacturer’s protocols.

### Microscopy

Airyscan images were taken with a Plan-Apochromat 63x/1.4 Oil DIC M27 objective with a 152 µm sized pinhole with master gain 800 using a Zeiss LSM 880 confocal microscopy equipped with an Airyscan detector (Carl Zeiss). 4 laser lines at 405, 488, 561, and 633 nm and a filter set for each line were used for taking 20-25 series of z-plane Airyscan confocal images with a step of 0.185µm or 0.247 µm for each channel. Pixel dwelling time was 2.33 µs and x and y step sizes were 43nm. 3D-Airyscan processing was performed with the Zen Black 2.3 SP1 program. For some images, Z-plane linear transitional alignment was done by using the Zen Blue 2.5 program. Contrast of images for each fluorophores channel was adjusted based on FMO (Fluorescence minus one) control samples that were prepared and taken on the same day of each experiments. To visualize cell fragments, sorted CD3+CD14mid cells were immobilized using CyGel Sustain (Abcam) according to manufacturer recommendations. Three dimensional rendering of cellular fragments (**Figure 2D**) was created in Imaris 9.1 software (Bitplane).

### Bulk memory CD4+ T cell sorting

Frozen PBMC were thawed and stained with fixable viability dye eFluor506 (eBiosciences) and various combinations of the antibodies listed in **Table supplement 1** as described in the flow cytometry section above. Memory CD4 T cell sorting (see gating strategy **Figure supplement 1A**) was performed on a BD Aria III/Fusion cell sorter (BD Biosciences). 100,000 memory CD4+ T cells were sorted into TRIzol LS reagent (Invitrogen) for RNA extraction.

### RNA sequencing and analysis

RNA sequencing and analysis of memory CD4+ T cells from LTBI infected subjects was performed as described in (Picelli et al., 2013; Seumois et al., 2016) and quantified by qPCR as described previously (Seumois et al., 2012). 5 ng of purified total RNA was used for poly(A) mRNA selection, full length reverse-transcription and amplified for 17 cycles, following the smart-seq2 protocol (Picelli et al., 2013; Seumois et al., 2016). After purification with Ampure XP beads (Ratio 0.8:1, Beckmann Coulter) and quantification (Picogreen assay, Invitrogen), 1ng of cDNA was used to prepare a Nextera XT sequencing library with the Nextera XT DNA library preparation and index kits (Illumina). Samples were pooled and sequenced using the HiSeq2500 (Illumina) to obtain at least 12 million 50-bp single-end reads per library. The single-end reads that passed Illumina filters were filtered for reads aligning to tRNA, rRNA, and Illumina adapter sequences. The reads were then aligned to UCSC hg19 reference genome using TopHat (v 1.4.1) (Trapnell, Pachter, & Salzberg, 2009), filtered for low complexity reads, and parsed with SAMtools (Li et al., 2009). Read counts to each genomic feature were obtained using HTSeq-count program (v 0.6.0) (Anders, Pyl, & Huber, 2015) using the “union” option. Raw counts were then imported to R/Bioconductor package DESeq2 (Love, Huber, & Anders, 2014) to identify differentially expressed genes among samples.

## Acknowledgments

We thank Dr Chery Kim and all present and past members at the flow cytometry core facility at the La Jolla Institute for Immunology for assistance in cell sorting and technical discussion. We thank Dr. Zbigniew Mikulski from the microscopy core at the La Jolla Institute for Immunology for assistance and technical advice on microscopy imaging. We thank Yoav Altman at the Sanford Burnham Prebys flow cytometry core for technical assistance with imaging flow cytometry. We thank Dr. Joe Trotter from the R&D Advanced Technology Group at BD Biosciences for useful technical discussion about cytometry instrument fluidic systems. Research reported in this manuscript was supported by the National Institute of Allergy and Infectious Diseases division of the National Institutes of Health under award number U19AI118626, R01AI137681, HHSN272201400045C, S10OD021831 and S10OD016262. The content is solely the responsibility of the authors and does not necessarily represent the official views of the National Institutes of Health. Imaging flow cytometry was supported by the James B. Pendleton Charitable trust.

## Authorship and conflict-of-interest statements

conceptualization: J.G.B, M.P, C.L.A, M.B, A.S and B.P; investigation: J.G.B, M.P, C.L.A, D.W, R.d.S.A, Y.J, V.S and G.S; resources: C.L.A, J.A.G, S.P, G.P, A.W, D.V, B.G, R.T, A.D.S, R.H, M.S, R.T, K.L and P.V; writing (original draft preparation): J.G.B and B.P; writing (review and editing): all authors; funding acquisition: A.S and B.P. All authors declare no competing interests.

## Data deposition

Sequencing data is accessible online through Gene Expression Omnibus (accession numbers GSE84445 and GSE99373, https://www.ncbi.nlm.nih.gov/geo) and Immport (Study number SDY820, http://www.immport.org). All other data is available in the main text or the supplementary materials.

---

**Figure 1 – figure supplement 1.**
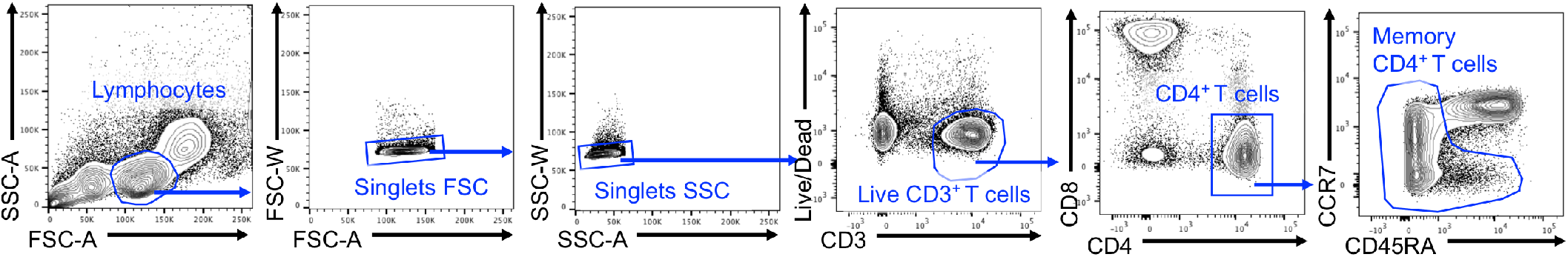
Gating strategy to isolate bulk memory CD4+ T cells.

**Figure 1 – figure supplement 2.**
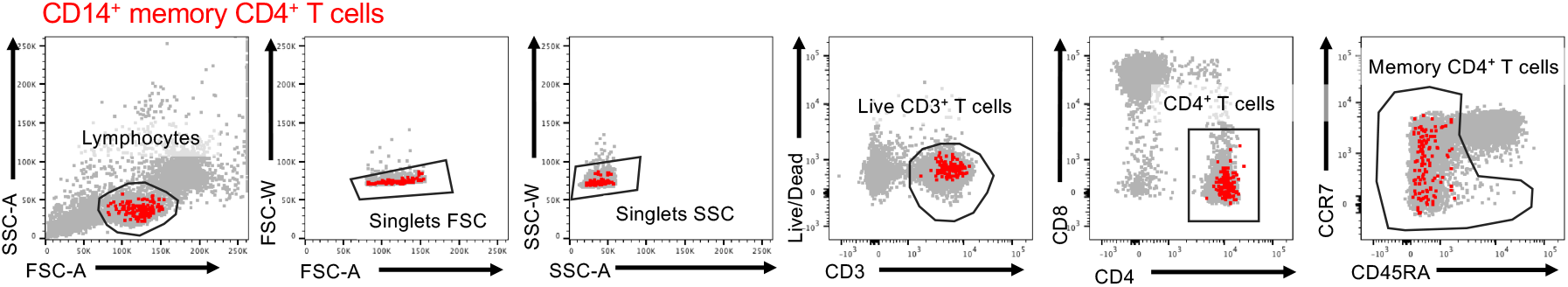
Backgating of CD14+ cells within sorted memory CD4+ T cells.

**Figure 2 – figure supplement 1.**
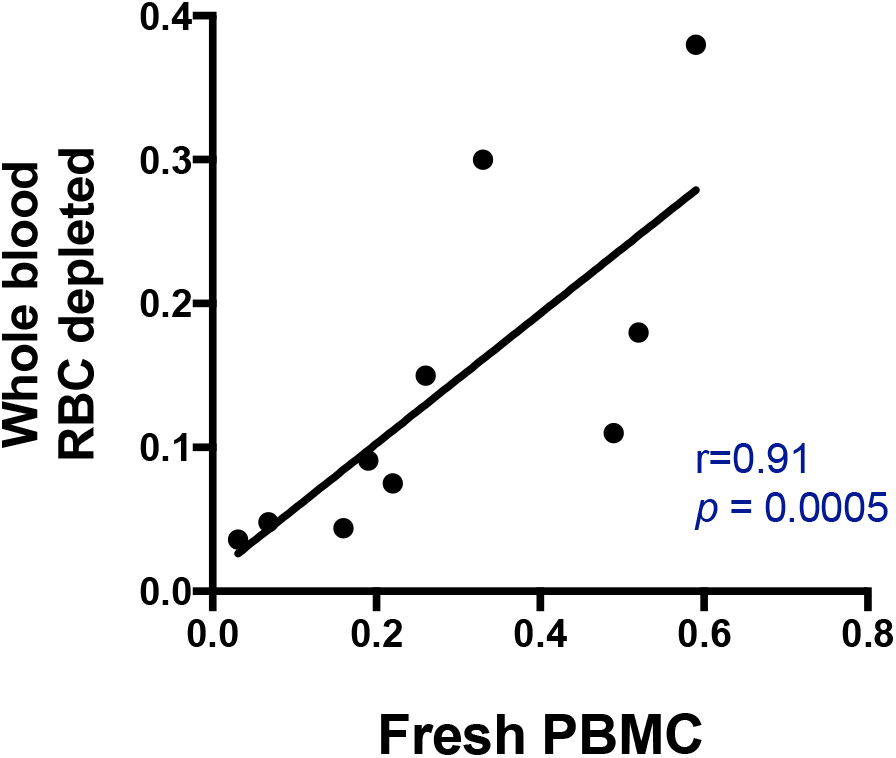
Non-parametric spearman correlation between CD3+CD14+ frequencies in whole blood versus fresh PBMC. Red blood cells were magnetically depleted from fresh whole blood using the EasySep RBC depletion kit (STEMCELL technologies) according to the manufacturer’s instructions. Data was derived from 10 independent blood draws of healthy subjects.

**Figure 2 – figure supplement 2.**
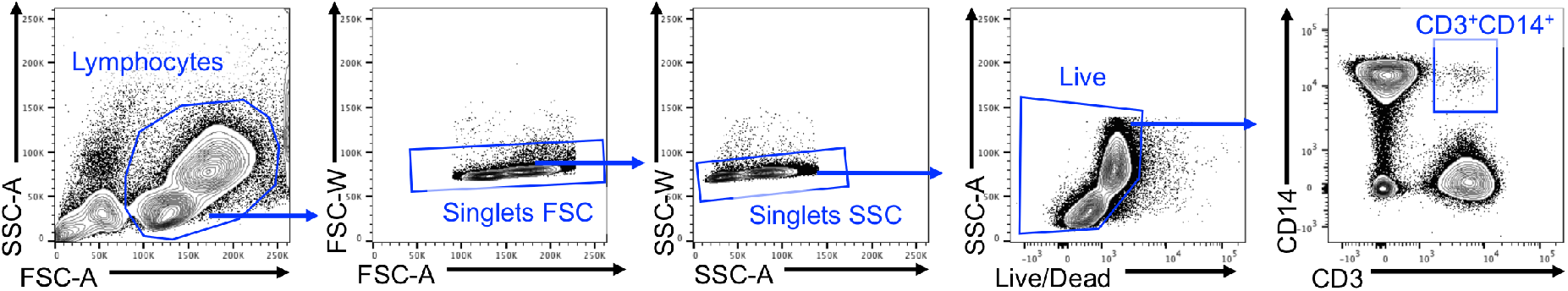
Gating strategy to isolate CD3+CD14+ cells.

**Figure 2 – figure supplement 3.**
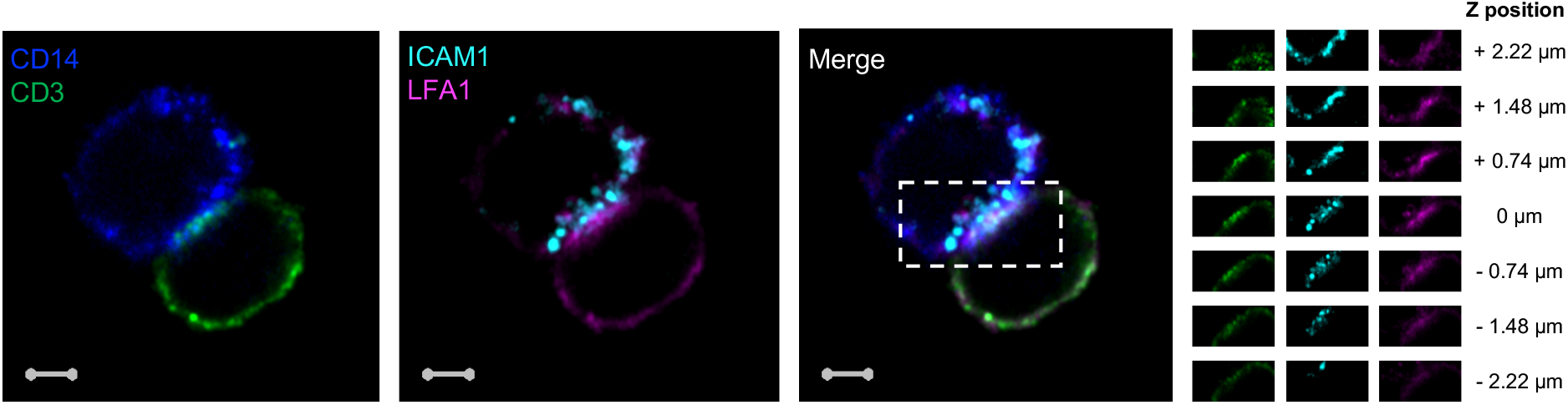
Accumulation of CD3, LFA1 and ICAM1 at the interface of a T cell:monocyte complex. Single z-plan (0µm) images (*left*) and z-plane stacks (*right*) of the region marked (dashed rectangle) from one sorted CD3+CD14+ Tcell:monocyte complex displaying accumulation of LFA1 and ICAM1 at the interface. Images show expression of CD14 (blue), CD3 (green), ICAM1(Cyan), and LFA1 (Magenta). Relative z-positions are indicated on the right, and scale bars represent 2 µm.

**Figure 3 – figure supplement 1.**
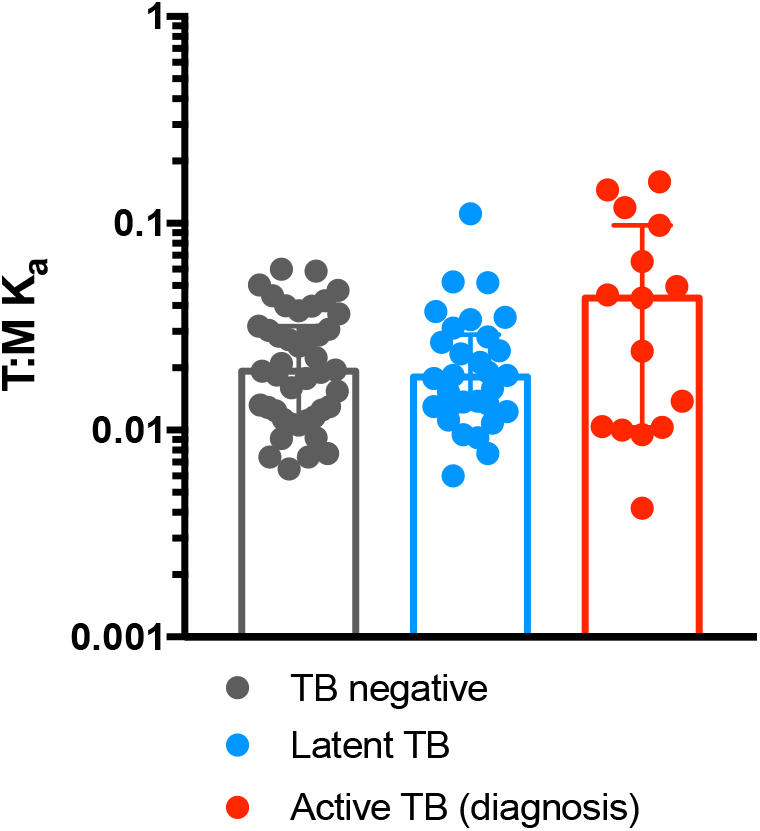
T cell:monocyte constant of association Ka in subjects with active TB, latent TB or TB uninfected individuals. T cell:monocyte constant of association Ka for was calculated as explained in Fig. 3B from active TB samples (n=15) collected at diagnosis from Sri Lanka, latent TB samples collected from subjects living in San Diego (n=22) or Peru (n=8), and TB uninfected samples collected from subjects living in San Diego (n=29) or Sri Lanka (n=14). Plots represent individual data points, median and interquartile range across all subjects within each cohort.

**Figure 3 – figure supplement 2.**
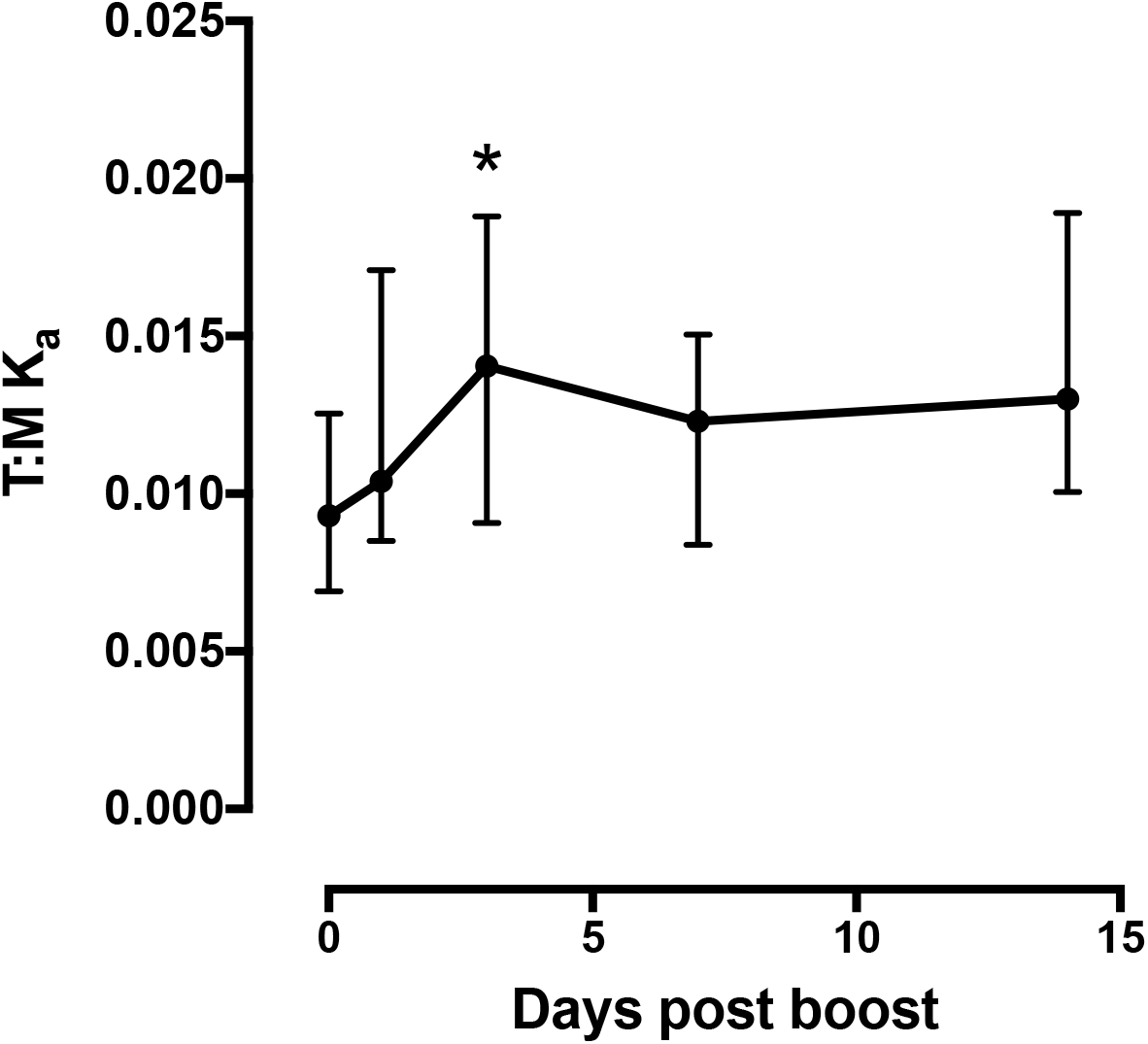
T cell:monocyte constant of association Ka fluctuates as a function of time following Tdap boost administration. Previously vaccinated healthy subjects (n=16) were re-immunized with Tdap and blood collected before, one day, three days, seven days and fourteen days post boost. Plots represent the median and interquartile range across all 16 subjects. T cell:monocyte constant of association Ka was calculated as explained in Fig. 3B.

**Figure 3 – figure supplement 3.**
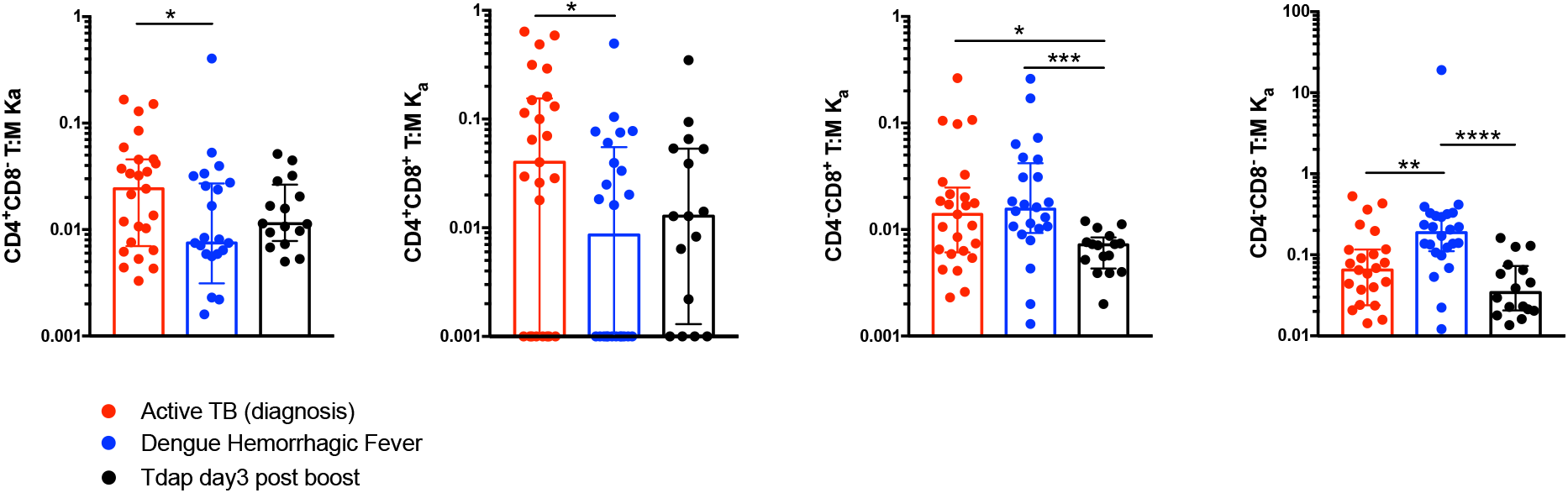
Comparison of constant of association Ka between monocytes and T cell subsets across different immune perturbations. Constant of association Ka for each T cell subset and monocytes was calculated as explained in Fig. 3B from active TB subjects at diagnosis (n=25), individuals with acute dengue hemorrhagic fever (n=24) and previously vaccinated healthy adults three day post boost with Tdap vaccine (n=16). Plots represent individual data points, median and interquartile range across all subjects.

**Figure 3 – figure supplement 4.**
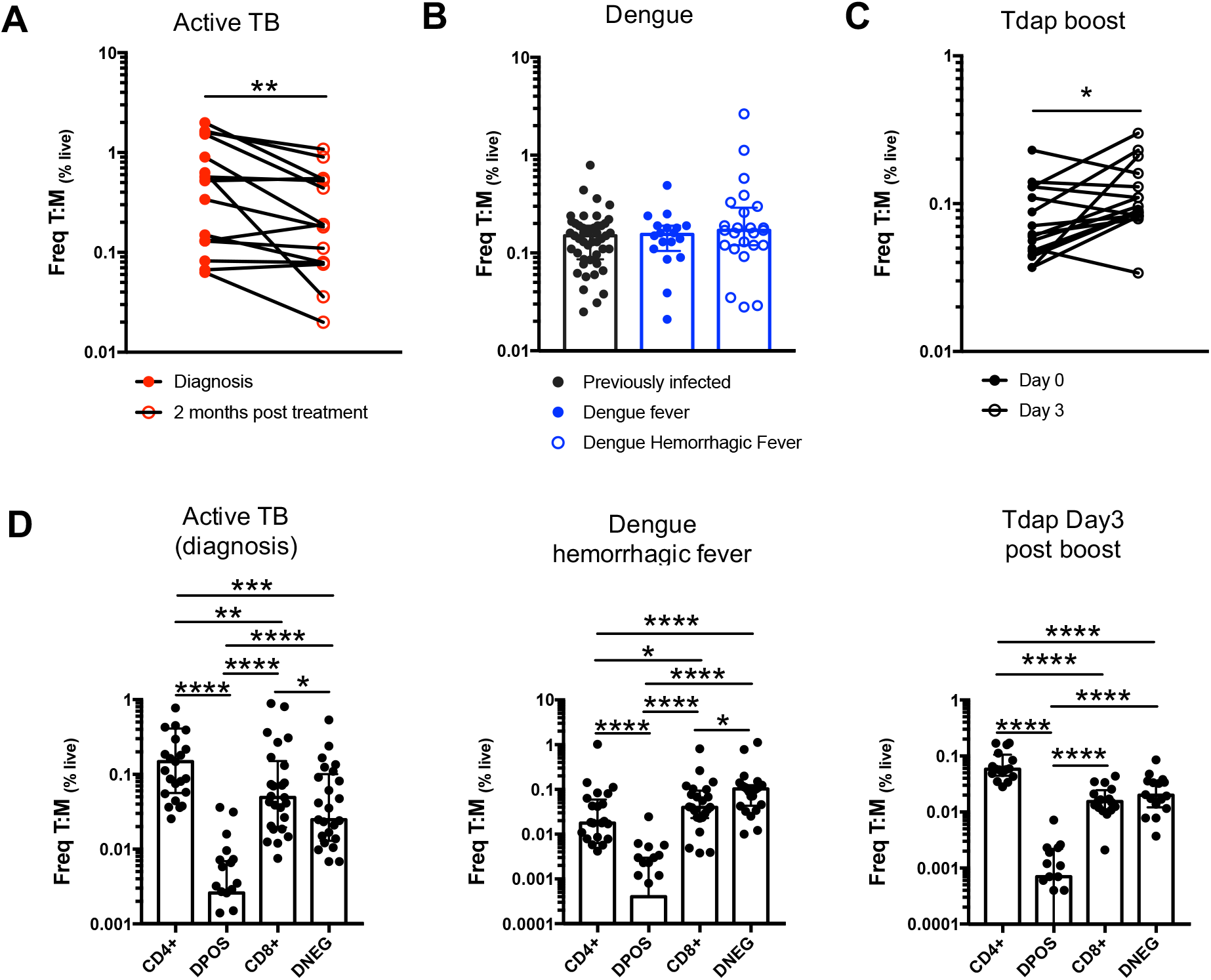
Frequencies of T cell:monocyte complexes in different immune perturbation models. Frequencies of T cell:monocyte complexes (and T cell subsets:monocyte complexes) expressed as percent of live cells were determined in active TB subjects at diagnosis (n=25) and two months post treatment (n=15), individuals with acute dengue hemorrhagic fever (n=24) and previously vaccinated healthy adults three days post boost with Tdap vaccine (n=16). Statistical differences over time and across cell populations within subjects were determined using the non-parametric paired Wilcoxon test; other statistical differences were determined using the non-parametric Mann-Whitney test; *, *p* < 0.05; **, *p* < 0.01; ***, *p* < 0.01; ****, *p* < 0.0001. Plots represent individuals data points, median and interquartile range across all subjects within each cohort.

